# Head-to-head comparison of CCN4, DNMT3A, PTPN11, and SPARC as suppressors of anti-tumor immunity

**DOI:** 10.1101/2022.04.01.486749

**Authors:** Anika C. Pirkey, Wentao Deng, Danielle Norman, Atefeh Razazan, David J. Klinke

## Abstract

**Background:** Effective communication between innate and adaptive immunity is essential for mounting an effective antitumor immune response. Emergent cancer cells likely secrete factors that inhibit this communication. To identify such factors, we applied an in vitro workflow that coupled a functional assay with proteomics to a syngeneic mouse model of melanoma known to be resistant to immunotherapies. Collectively, these in vitro results suggested CCN4, DNMT3A, PTPN11, and SPARC as secreted factors that potentially mediate immunosuppression and could potentially become easily accessible targets for novel immunotherapies. The objective of this study was to test for consistent clinical correlates in existing human data and to verify in vivo whether knocking out tumor cell production of these secreted factors improved immune-mediated control of tumor growth.

**Results:** Analyses of available human data indicate that high CCN4 expression is associated with reduced survival in primary melanoma patients. High DNMT3A, PTPN11, and SPARC expression showed no association with overall survival. In addition, scRNAseq data from patients with melanoma confirmed CCN4 is expressed by the tumor cells, as opposed to other cells in the tumor microenvironment. An experimental system was created using a CRISPR/Cas9 approach to generate knock out cell lines for each of the four genes of interest using the B16F0 mouse melanoma cell line. In a immunocompetent C57BL/6, DNMT3A, PTPN11, and SPARC knockouts had no effect on overall survival compared to mice challenged with wildtype B16F0 cells while the CCN4 knockout significantly increased survival. This increase in overall survival was lost when the CCN4 knockout cells were injected into severely immunocompromised NSG hosts, indicating that CCN4’s effect on the tumor microenvironment is immune mediated. A kinetic analysis leveraging a Markov Chain Monte Carlo approach quantified the various knockouts’ effect on cells’ intrinsic growth rate. The analysis shows CCN4 is the only knockout tested that decreased the net proliferation rate in immunocompetent mice compared to wildtype cells.

**Conclusions:** The results suggest that CCN4 is a mediator of immunosuppression in the melanoma tumor microenvironment and a potential collateral immunotherapy target.

## Background

The anti-tumor response achieved by immune checkpoint inhibitors is dependent not only on the presence of CD8+ T cells within the tumor (1), but on the successful coordination of response from a variety of cells and the secreted signals they use to communicate within the tumor microenvironment (2). For instance, dendritic cell release of IL-12 within the tumor microenvironment underpins the anti-tumor response to immune checkpoint inhibition (3). If this tonic IL-12 signaling is important for anti-tumor immunity, interrupting this intercellular signal should provide a survival advantage for malignant cells that arise during oncogenesis.

Motivated by the importance of tonic IL12 signaling and the evolutionary nature of oncogenesis, we previously asked whether tumor cells secrete factors that promote tolerance by suppressing immune function. In vitro co-culture models were used to test the importance of cytokines such as IL12 in eliciting an anti-tumor T cell response by examining whether the presence of a malignant cell interfered with the T-cell response to IL-12 (4). Using unbiased omics-based approaches to identify tumor-secreted factors that underpin cellular cross-talk, DNA methyltransferase 3 (DNMT3A) and Protein Tyrosine Phosphatase Non-Receptor Type 11 (PTPN11), which are enriched in tumor secreted vesicles, as well as the secreted proteins Cellular Communication Network Factor 4 or Wnt-inducible signaling pathway protein-1 (WISP1/CCN4) and Secreted Protein Acidic and Rich in Cysteine (SPARC) were highlighted as potential mediators of immunosuppression (4–6). While this prior work highlighted individual genes in a limited cellular context, head-to-head comparisons among these potential targets using an in vivo model and a CRISPR-based gene knock-out approach would be helpful for prioritization. Because of the inherent limitations of using mouse cells to attempt to capture the same pathways present in humans, it would also be beneficial to use a bioinformatics approach to test for consistent clinical correlates in existing data.

## Materials and Methods

### Cell lines and mouse models

The B16F0 (RDID: CVCL_0604), mouse melanoma cell line was obtained from American Tissue Culture Collection (ATCC, Manassas, VA). Upon receipt, cells were aliquoted and frozen in liquid nitrogen. For specific experiments, cell lines were brought out of freeze back, used within 10-15 passages, and routinely tested for mycoplasma contamination by PCR. When cultured, the cells were kept at 37°C in 5% CO2 in high-glucose DMEM (Cellgro/Corning, NY) supplemented with 10% heat-inactivated fetal bovine serum (Hyclone, UT), penicillin-streptomycin (Gibco) and L-glutamine (Lonza). Variants of the parental B16F0 cell line were generated by knocking out DNMT3A (DNMT3A-KO), PTPN11 (PTPN11-KO), SPARC (SPARC-KO) and WISP1/CCN4 (CCN4-KO) using a double nickase-based CRISPR/Cas9 approach as previously described (7). Plasmids were purchased from Santa Cruz Biotechnology, Inc. (Dallas, TX) and, following puromycin selection, selected colonies were expanded on 6-well plates, aliquoted and frozen in liquid nitrogen.

Female C57BL/6Ncrl and male NOD-scid IL2R *γ*^*null*^ immunodeficient mice (NSG), all six to eight weeks old, were purchased from Charles River Laboratories and The Jackson Laboratory, respectively. Upon receipt, animals were labeled and randomly assigned to treatment arms/cages, with a density of five mice per cage. All procedures involving animals were approved by the West Virginia University (WVU) Institutional Animal Care and Use Committee and performed at the WVU Animal Facility ((IACUC Protocol 1604002138).

### Western blot analysis

Western blotting was performed as previously described (7). Mouse anti-*β*-actin (C4), rabbit anti-WISP-1 (H-55, sc-25441) and rabbit antiSH-PTP2/PTPN11 (C-18, sc-280) were from Santa Cruz Biotechnology. Rabbit monoclonal antibodies, antiDNMT3A (D23G1) and anti-SPARC (D10F10), were from Cell Signaling Technology (Danvers, MA).

### In vivo tumor studies

C57BL/6Ncrl and NSG mice were each divided into five groups (n = 8 in C57BL/6 CCN4 WT and KO groups, else n = 5). 2.2 × 10^5^ B16F0 mouse melanoma cells or corresponding knockout varients in 100 *µ*L PBS were injected subcutaneously in the right flank of each mouse. Mice were monitored every day until the predefined endpoints. Two orthogonal diameters (width and length) were measured with a digital caliper. Tumor volume was calculated according to the formula 0.5236 x width^2^ x length.

### Statistical analysis

For the survival analysis, gene expression data and matching clinical profiles for patients diagnosed with primary melanoma (stages I and II) were downloaded from the skin cutaneous melanoma (SKCM) arm of the Cancer Genome Atlas (TCGA) database (accessed on 11/25/2019) using the “TCGAbiolinks” (V2.14.1) package in R (V3.6.3). Kaplan-Meier survival curves were created using the Surv() and survfit() functions in the “survival” package (V3.1.8) and then plotted using the ggsurvplot() function in the “survminer” package (V0.4.6). The Cox proportional hazards model was created using the analyse_multivariate() and forest_plot() functions in the “survival Analysis” package (V0.1.3)

Single-cell RNA sequencing (scRNAseq) data from tumor samples obtained from patients diagnosed with primary melanoma were downloaded from Gene Expression Omnibus (accession numbers GSE115978 and GSE72056) (8, 9). Only cells classified as malignant were included for this analysis. For each gene of interest (DNMT3A, PTPN11, SPARC, WISP1/CCN4), non-zero counts in respective gene expression indicated a “gene-positive” tumor cell. A binomial test with a null proportion of 0.5% was used to determine samples with a significant proportion of cells expressing each gene of interest. The resulting p-value is the probability of observing a proportion greater than or equal to the sample data in a population where 1 in 200 cells will express the gene of interest.

Fisher’s exact test determined whether the proportion of gene-positive samples in the bulk TCGA patient data is comparable to that in the single-cell tumor samples. A p-value greater than 0.05 favors the null hypothesis that any difference in the proportion of cells expressing the gene of interest in the single-cell versus bulk sequencing data is explained by random chance. A p-value less than 0.05 indicates the two proportions are not equal.

### Markov Chain Monte Carlo analysis

The Markov Chain Monte Carlo (MCMC) analysis was completed in R (V3.6.3) as previously described (10). In brief, longitudinal tumor volume data was collected using caliper measurements was linearly transformed using a log10 scale. A function in R was created to complete the MCMC analysis for a single set of starting parameters and a single data set (see MHmcmc function in GitHub repository). The MCMC analysis uses a Metropolis Hastings algorithm and assumes a uniform prior for the model parameters. Chains were generated using 100,000 steps and a 20% acceptance fraction of proposed steps. The analysis was run three times for each dataset using different starting points to ensure convergence was independent of the starting point. The Gelman Rubin Convergence Diagnostic was used to confirm that the variance between the chains was smaller than the variance within the chains. A value of the Gelman-Rubin statistic close to 1.0 indicates that each chain converges to the same solution regardless of starting point. Following discarding the first 1,000 steps as a burn-in, the chains were thinned to 1,000 data points and, assuming convergence, represent samples from the posterior distributions in the model parameters.

## Data Availability

The key datasets used in the analysis can be obtained from the following sources:

- Bulk tissue transcriptomics profiling data using Illumina RNA sequencing was accessed from the SKCM arm of the Cancer Genome Atlas. Data were downloaded from TCGA data commons using the “TCGAbiolinks” (V2.14.1) package in R (V3.6.3). Gene expression data were expressed in counts.
- The single cell RNAseq datasets used in the analysis for this article are available in Gene Expression Omnibus repository with the following GEO accession numbers: GSE115978 and GSE72056.

## Code Availability

The code used in the Markov Chain Monte Carlo analysis can be obtained from the following GitHub repository:

- https://github.com/arcoolbaugh/B16-In-Vivo-Screen

## Results

### CCN4 is associated with reduced survival in primary melanoma patients

To assess the clinical context of DNTM3A, PTPN11, SPARC, and CCN4 in primary melanoma patients, complete survival histories of 95 patients diagnosed with primary melanoma were obtained from the SKCM arm of TCGA for statistical analysis. The patients were stratified into two groups based on gene expression for the gene of interest and their overall survival was visualized using Kaplan-Meier survival curves. Considered individually, expression of DNMT3A (Fig. 1A), PTPN11 (Fig. 1B), and SPARC (Fig. 1C) were not statistically significant correlates of overall survival (p-values = 0.14, 0.12, and 0.11 respectively). In contrast, increased expression of WISP1/CCN4 (Fig. 1D) independently correlated with reduced overall survival of patients diagnosed with primary melanoma (p-value = 0.0014).

**Figure 1.**
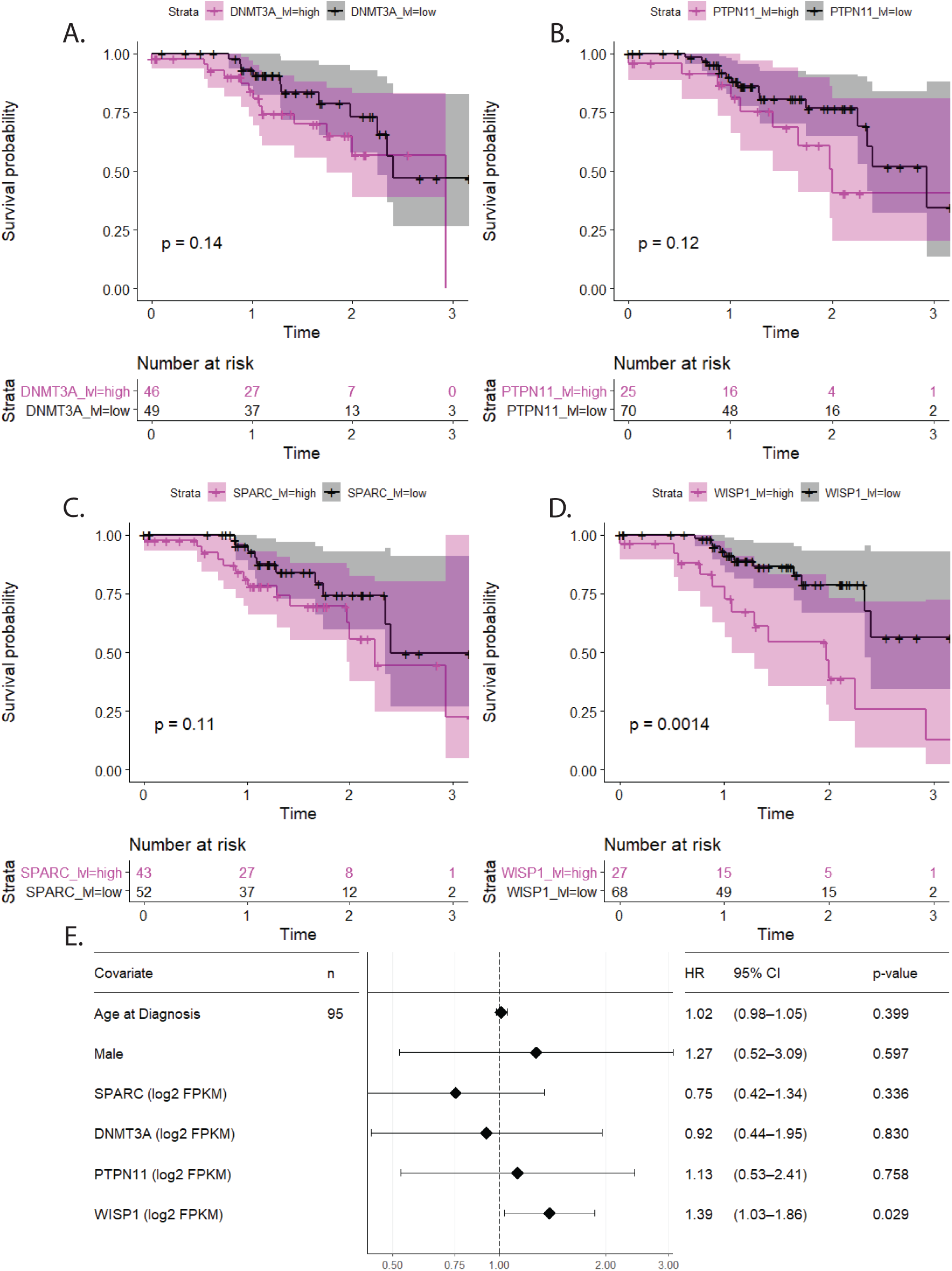
CCN4 expression is associated with reduced overall survival of patients diagnosed with primary melanoma. (A-D) Kaplan-Meier estimate of overall survival of primary melanoma patients stratified by (A) DNMT3A, (B) PTPN11, (C) SPARC, and (D) WISP1/CCN4 transcript abundance. Sample numbers are given in the table below the corresponding curve. P-values indicated in the plot area were calculated using the Peto and Peto modification of the Gehan-Wilcoxon test. (E) Graphical summary of the multivariate Cox proportional hazards model for overall survival based on the population characteristics age, sex, and expression of DNMT3A, PTPN11, SPARC, AND WISP1/CCN4. TCGA dataset accessed on 11/25/2019.

While common covariates may be lurking variables that underpin these independent test results, expression of DNMT3A, PTPN11, SPARC, and CCN4, along with age at diagnosis, gender, and tumor stage were used as covariates in a multivariate Cox proportional hazards model to assess their statistical association with overall survival. Of all covariates considered, increased expression of CCN4 was the only covariate found to be statistically significant (HR = 1.36, p-value = 0.032, see Fig. 1E).

### CCN4 is expressed by tumor cells in a subset of primary melanoma patients

To verify that the genes of interest were being produced by the tumor cells as opposed to other cells present in the bulk RNAseq data from TCGA, scRNAseq data of malignant cells from patients diagnosed with primary melanoma were analyzed for the proportion of cells in each sample that expressed the desired gene. While nearly all samples showed statistically significant expression of DNMT3A, PTPN11, and SPARC (see Fig. 2 A-C), only nine of the twenty-six samples showed statistically significant expression of CCN4 (Fig 2 D). A Fisher’s exact test was used to test whether the proportion of treatment naïve samples expressing CCN4 was equal to the proportion of high CCN4 expressing samples in the TCGA sample set (p-value = 1 indicating the proportions are not statistically different). These scRNAseq data combined with the TCGA data indicates that malignant melanocytes are a source of CCN4 within the tumor microenvironment. To determine whether CCN4 affects tumor progression by modulating anti-tumor immunity, we turned to in vivo mouse models.

**Figure 2.**
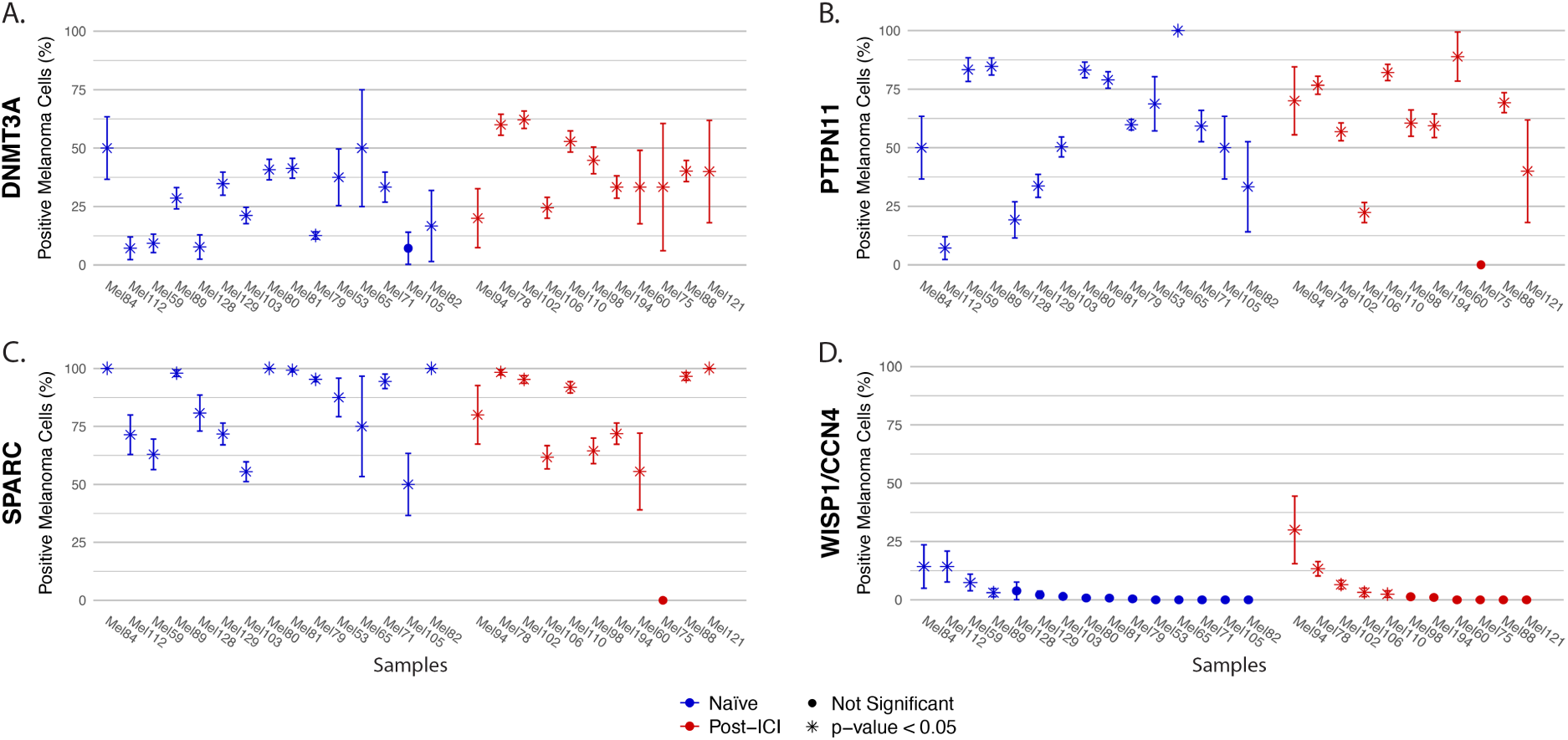
An analysis of publicly available scRNAseq data shows that WISP1/CCN4 is expressed in a subset of the tumors of primary melanoma patients. The proportion of melanoma cells expressing (A) DNMT3A, (B) PTPN11, (C) SPARC, and (D) WISP1/CCN4 in each tumor sample. Error bars represent the standard error of the sample proportion, given a binomial distribution. Samples are separated into treatment-naïve (blue) and post-immune checkpoint inhibitor (post-ICI) therapy (red). Asterisks (*) mark samples with a significant proportion of cells expressing the gene of interest, determined by a binomial test with a null frequency of 0.5%.

### Knockout of CCN4 in immunocompetent mice enhances overall survival

To develop an experimental system, a double nickase-based CRISPR/Cas9 approach was used to generate knock out cell lines for each of the four genes of interest using the syngeneic B16F0 melanoma cell line (11). Knockout was verified by western blot (Supplemental Figure 1). Wildtype (WT) B16F0 and each knockout (KO) variant were used to challenge both C57BL/6 and NSG mouse strains by subcutaneous injection, where overall survival and dynamic tumor growth were recorded as outcome metrics. In vivo challenge using syngeneic C57BL/6 and NSG mouse strains were used to model tumor growth in the presence or near complete absence, respectively, of host immunity. In immunocompromised NSG mice, DNMT3A, PTPN11, SPARC, and CCN4 (Fig. 3 A-D, left panels) all had no statistically significant impact on overall survival (n = 5 mice/group, p-value = 1, 0.3, 1, and 0.3 respectively). Similar to our observations using the primary melanoma patient data, in C57BL/6 mice receiving DNMT3A KO, PTPN11 KO, and SPARC KO cells (Fig. 3 A-C, right panels) showed no statistically significant difference in survival compared with mice challenged with WT B16F0 cells (n = 5 mice/group, p-value = 0.8, 0.3, and 0.8 respectively). Knocking out CCN4, however, led to a statistically significant increase in overall survival in C57BL/6 mice receiving CCN4 KO cells (Fig. 3D, right panel) (n = 8 mice/group, p-value = 0.01).

**Figure 3.**
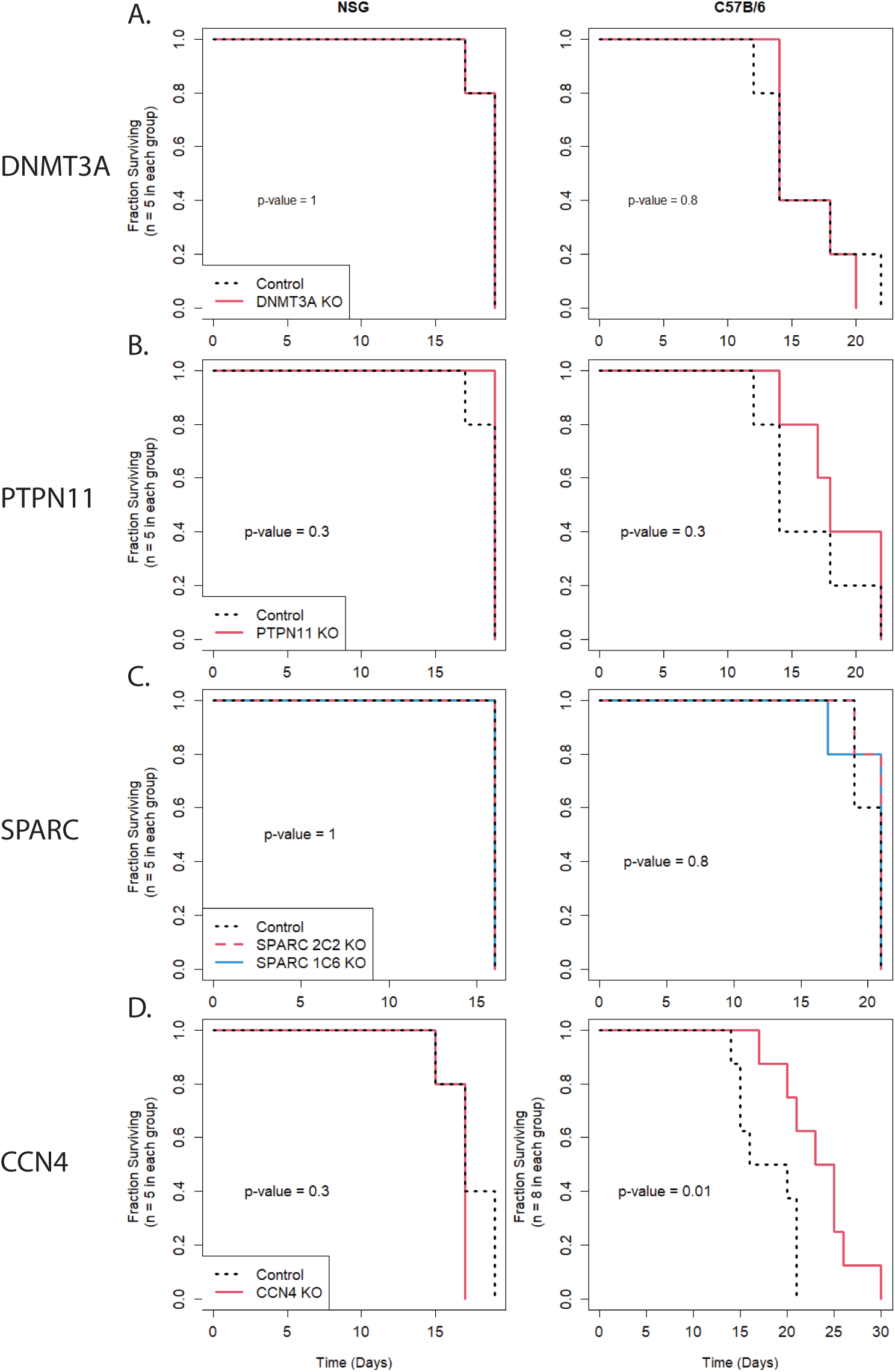
CCN4 knock out increases overall survival in immunocompetent mice. (A-D) Kaplan-Meyer survival analysis of four knock out variants of the B16F0 melanoma model. DNMT3A (A), PTPN11 (B), SPARC (C), and WISP1/CCN4 (D) knockout cells were injected into NSG (left panel) and C57B/6 (right panel) mice to compare survival against the wild type B16F0 cell line. Control group is indicated by the black dotted line, while the knockout groups are indicated in red. Two variants of SPARC knockout were tested, where the 2C2 knockout is represented by the red dashed line and the 1C6 knockout is represented by the blue solid line. The n for each group is 5, except the WISP1/CCN4 injected in C57B/6 mice, where n = 8 for each group. P-values indicated were calculated using the Peto and Peto modification of the Gehan-Wilcoxon test.

### Knockout of CCN4 increases growth rate of B16F0 melanoma tumors

Comparing tumor burden at a single time point is often used as an endpoint in mouse model experiments. However, comparisons at a single time point can be influenced by latent variables associated with by the experiemental process, such as differences in intrinsic growth rate or in immunoediting introduced by the gene editing process or differences in the initial bolus of tumor cells injected that survive to form tumors (12). To better quantify the impact that knocking out the genes of interest have on tumor burden, we recorded tumor growth trajectories in each NSG and C57BL/6 mouse once the tumors were palpable (Day 7) until the terminal endpoint was reached. In vivo growth profiles were compared quantitatively using a mathematical model of the exponential tumor growth. The model assumes that the overall growth rate of the tumor at any point in time depends on the number of tumor cells present, the rate constant associated with intrinsic proliferation (kp), which accounts for both cell division and natural cell death due to mechanisms independent of host immunity; and the rate constant associated with immune-mediated death of the cells (*k*_*D*_) that is only present in the immunocompetent C57BL/6 mouse, that is in NSG mice, *k*_*D*_ is equal to 0 (see Supplemental Figure 2). As summarized in the methods, a Bayesian approach based upon a Markov Chain Monte Carlo (MCMC) method was used to estimate the uncertainty of the model parameters, given the available tumor volume data. In estimating the posterior distributions in the model parameters, the initial tumor bolus (To) was estimated for each mouse while the net tumor growth rate parameter (*k*_*P*_ or 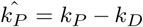) was shared among all mice of an experimental group (10, 13)

Representative traces of three independent MCMC chains and the corresponding Gelman Rubin statistics are shown in Supplemental Figure 3. Samples from the converged Markov Chains were used to generate posterior distributions in the model predictions. These were compared against experimental data and show that the error follows a normal distribution, suggesting that the mathematical model accurately captures the tumor growth trajectories in these mice.

To compare the posterior distributions of the model parameters, we visualized the log10 ratio of the net tumor growth rate parameter of the specific cell line variant in C57BL/6 mice 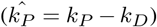 to the tumor growth rate parameter of the same cell line variant in NSG mice (*k*_*P*_) (Fig. 4 A-E, columns 3-4). Statistical differences in the posterior distributions of the log10 ratio of tumor growth rate parameters were assessed using a Pearson’s Chi-squared test, where comparisons among all WT versus KO groups were considered statistically significant (p-value < 0.05).

**Figure 4.**
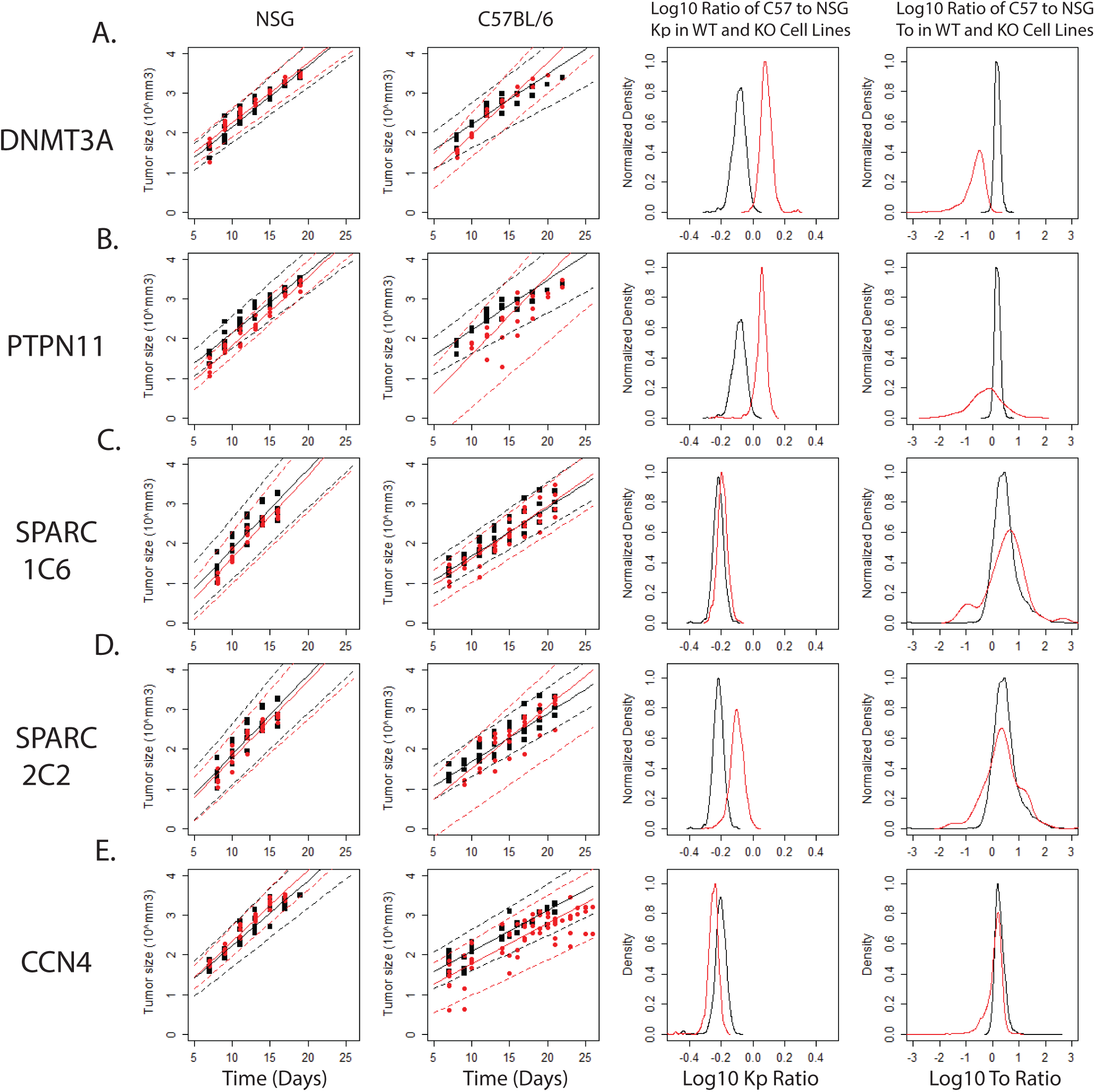
Knockout reduces tumor growth rate of B16F0 tumors in immunocompetent mice. CCN4 Knockout reduces tumor growth rate in immunocompetent mice. (A-E) B16F0 mouse melanoma with (A) DNMT3A, (B) PTPN11, (C) SPARC 1C6, (D) SPARC 2C2, or (E) WISP1/CCN4 knockout (red) and wildtype (black) cells were each injected into a cohort of NSG mice (panel 1, n=5 per group) and C57B/6 mice (panel 2, n=5 except for WISP1/CCN4 group n=8). The solid lines represent the average trendline for all data while the dotted lines enclose the 95% confidence interval. Markov Chain Monte Carlo analysis was used to identify a posterior distribution for the slope (logarithmic tumor growth rate, Kp) and intercept (logarithmic initial tumor injection size, To) of each experiment. Data is represented as the natural log of the ratio of the Kp (panel 3) or To (panel 4) in C57B/6 mice compared to NSG mice. Statistical significance was assessed using a Chi-squared test with simulated p-value, and all p-values were found to be < 1E-5 with the except of the shift in Ko of the SPARC 1C6 KO, which found p = 0.036.

In examining groups receiving WT cells (Fig. 4, black curves), the tumor growth rate parameter was consistently greater in NSG mice compared to C57BL/6 (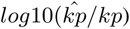, corresponding to a posterior log10 ratio distributed at values less than 0). In addition, injecting the same number of WT cells produced a slightly greater bolus of tumor-initiating cells in C57BL/6 compared to NSG mice, that is the distributions in log10 ratios were consistently centered in the positive domain. In considering KO variants (red curves in Fig 4), DNMT3A KO and PTPN11 KO variants produced a lower bolus of tumor initiating cells compared to their WT counterparts. Interestingly, the growth rate parameters for both DNMT3A KO and PTPN11 KO cells in C57BL/6 were higher than in NSG mice, that is the log10 ratio was distributed with values greater than 0 (see Fig. 4B). As *k*_*D*_ is defined as a positive definite number, this implies that the presence of immune cells enhances the intrinsic growth rate of DNMT3A KO and PTPTN11 KO cells. For SPARC and CCN4 KO variants, the initial bolus of tumor initiating cells were similar to WT. While the growth rate parameters for both SPARC and CCN4 KO variants were lower in C57BL/6 mice than NSG mice, the comparison between WT and SPARC KO variants suggests that the absence of SPARC did not increase the net propensity for cell death in the presence of host immunity. In contrast, CCN4 was the only knockout variant to show a decrease in 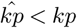 compared to the WT cells (Fig 4E). Given the similarity in initial boluses of tumor initiating cells upon CCN4 KO, the decrease in net growth rate suggests that immune-mediated killing of B16F0 is enhanced in the absence of tumor-derived CCN4, unlike that seen upon loss of tumor-derived SPARC, DNMT3A, and PTPN11.

## Discussion

This work explored four gene products and their potential in vivo to mediate immunosuppression in melanoma. DNMT3A is known to play a key role in the regulation of gene expression through the methylation of a set of genes necessary for late-stage embryonic development. Prior work implicated DNMT3A overexpression in tumorigenesis of liver cancer, colon cancer, and chronic myeloid leukemia, but the evidence and its mechanism of action remain unclear (14–16). Minimal evidence of the effect of DNMT3A exists in melanoma, where the only primary research using an in vivo model involved the use of an shRNA plasmid to knockdown DNMT3A in B16 cells (17). One of the limitations of shRNA is that it has off-target effects and transfecting in a shRNA plasmid can engage toll-like receptors eliciting an interferon response. While knocking down DNMT3A elicited an interferon response, the absence of control experiments, such as a rescue experiment with an RNAi-resistant target gene, undermine Deng and co-workers’ conclusions. PTPN11 is known to have a role in negatively regulating both IL-2 and T-cell receptor signaling pathways and is enriched in exosomes from the B16F0 melanoma cell line (5). This enrichment is able to suppress T-cell proliferation, which is typically associated with associated with increased tumor growth, decreased patient survival, and poor response to immunotherapies such as immune checkpoint inhibitors (5, 8). SPARC overexpression has been linked with the promotion of mesenchymal traits that enhance metastatic potential and ultimately immune evasion (18, 19). However, based on the available human data and our in vivo growth study, PTPN11 and SPARC appear to have no significant impact on tumor growth when compared in immunocompromised and immunocompetent backgrounds. Interestingly, our analysis indicates that the knockouts of DNMT3A and PTPN11 actually increase the intrinsic growth rate of B16F0 cells while knocking out SPARC had no effect. Being able to parse differences in intrinsic growth rates from changes in net tumor growth due to immunoediting is one of the advantages of this in vivo-in silico approach.

While this work has identified WISP1/CCN4 as enabling escape from immunoediting in the immunotherapy resistant B16F0 melanoma line, the mechanism of action was not identified here. Previous work has implicated WISP1/CCN4 as an actor in invasion and metastasis through its promotion of the epithelial-mesenchymal transition as well as immunosuppression in the tumor microenvironment, leading to increased tumor growth in an immunocompetent setting (7, 12). Similarly, promotion of metastasis and correlation with poor prognosis is seen in hepatocellular carcinoma and breast cancers (20, 21). Current work shows decreased infiltration of CD45+ cells and NK cells and increased granulocytic myeloid derived suppressor cells in tumors secreting CCN4 (22, 23).

Recent work to identify possible mediators of treatment resistance and immunosuppression that could be utilized as therapeutic targets has begun to rely on CRISPR/Cas9 screenings as their primary method (24, 25). However, while these screenings are high throughputin identifying potential therapeutic targets, they also identify intercellular proteins that may be difficult to target in therapy due to lack of access or that may carry increased risk for toxicity or adverse reactions based on their function in normal cells. Moreover, any screening approach has implicit bias in enriching for hits. CRISPR/Cas9 screens focus on identifying genes that alter intrinsic susceptibility to immunoediting. Phenotypic screens, such as an ability to suppress heterocellular crosstalk, coupled with unbiased proteomic methods, as described in Kulkarni et. al (4), coupled with in vivo validation, as described here, is an alterative. While the number of potential targets may be lower compared with a CRISPR/Cas9 approach, identifying secreted targets, like CCN4, hopefully has a clearer path forward for clinical translation.

## ACKNOWLEDGEMENTS

This work was supported by National Science Foundation (NSF CBET-1644932 to DJK) and National Cancer Institute (NCI 1R01CA193473 to DJK). The content is solely the responsibility of the authors and does not necessarily represent the official views of the NSF or NCI.

## AUTHOR CONTRIBUTIONS

These contributions follow the International Committee of Medical Journal Editors guidelines: http://www.icmje.org/recommendations/. Conceptualization: DJK; Study Design: DJK; Data Acquisition: DJK, WD, and AR; Data Analysis: ACP and DN; Data Interpretation: ACP and DJK; Funding acquisition: DJK; Methodology: DJK; Project administration: DJK; Software: DJK and ACP; Supervision: DJK; Writing - original draft: ACP, DN, and DJK; Writing - review & editing: all authors.

## COMPETING FINANCIAL INTERESTS

The authors declare no competing financial interests.

---

**Fig. S1.**
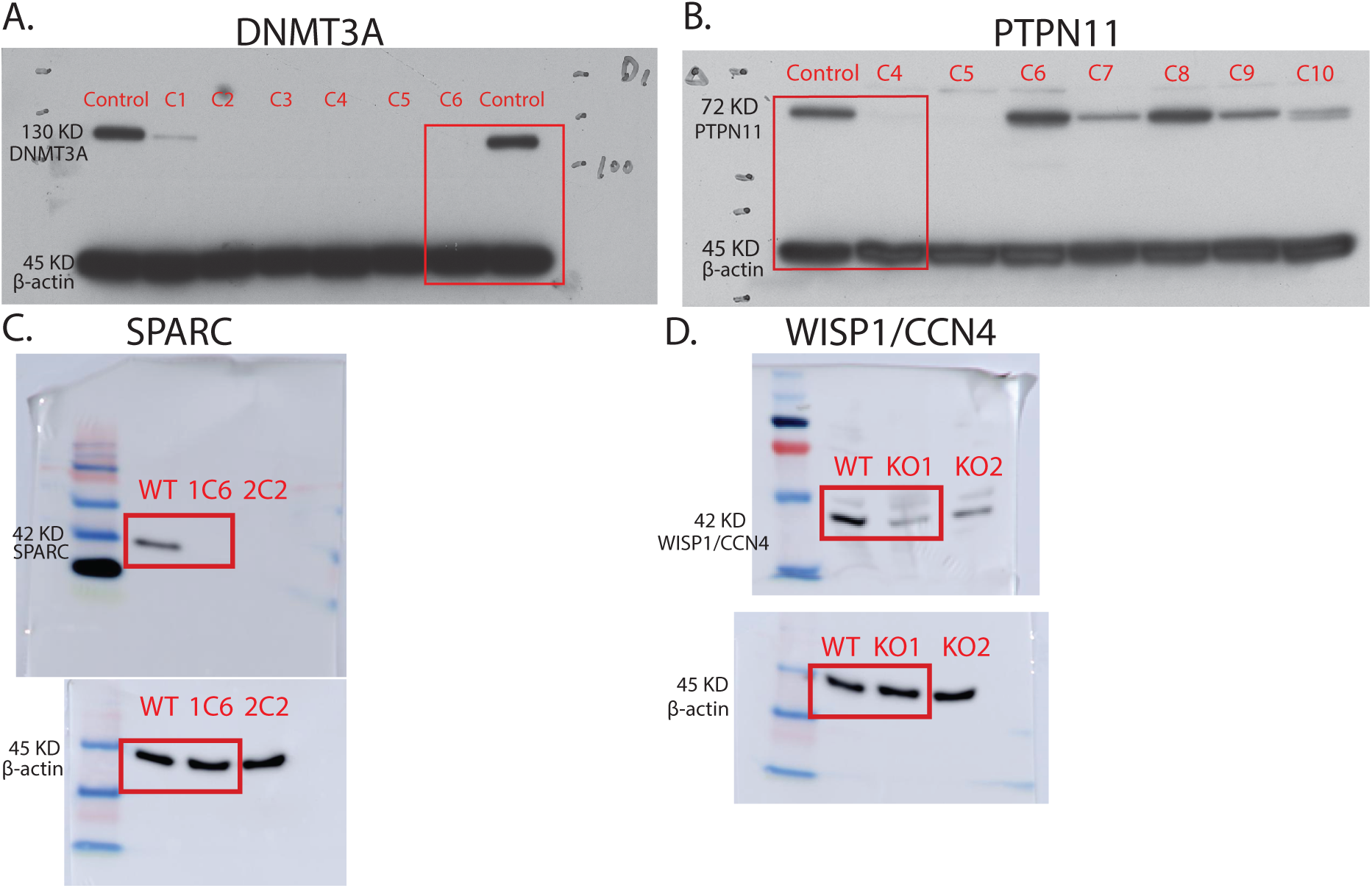
Western blots were used to confirm successful knockout of each gene of interest. Western blot images confirming the knockout of (A) DNMT3A, (B) PTPN11, (C) SPARC, and (D) WISP1/CCN4 in B16F0 cells.

**Fig. S2.**
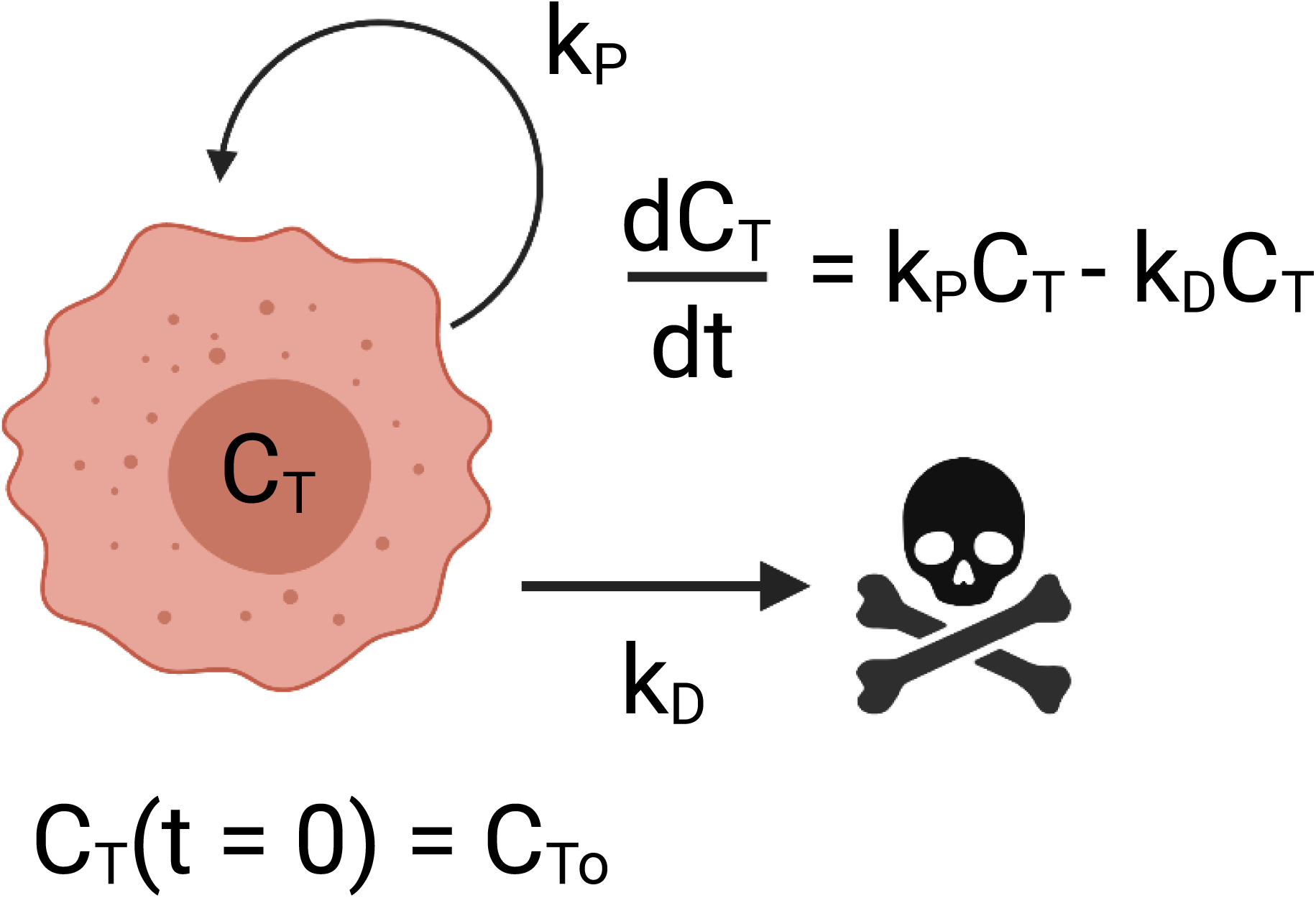
A schematic of the tumor growth model assumed in the Markov Chain Monte Carlo Analysis. The model accounts for an intrinsic net cell growth rate (*k*_*P*_) that encompasses cell division and non-immune mediated cell death while any immune mediated death that takes place in the C57BL/6 mice is accounted for using a separate constant (*k*_*D*_).

**Fig. S3.**
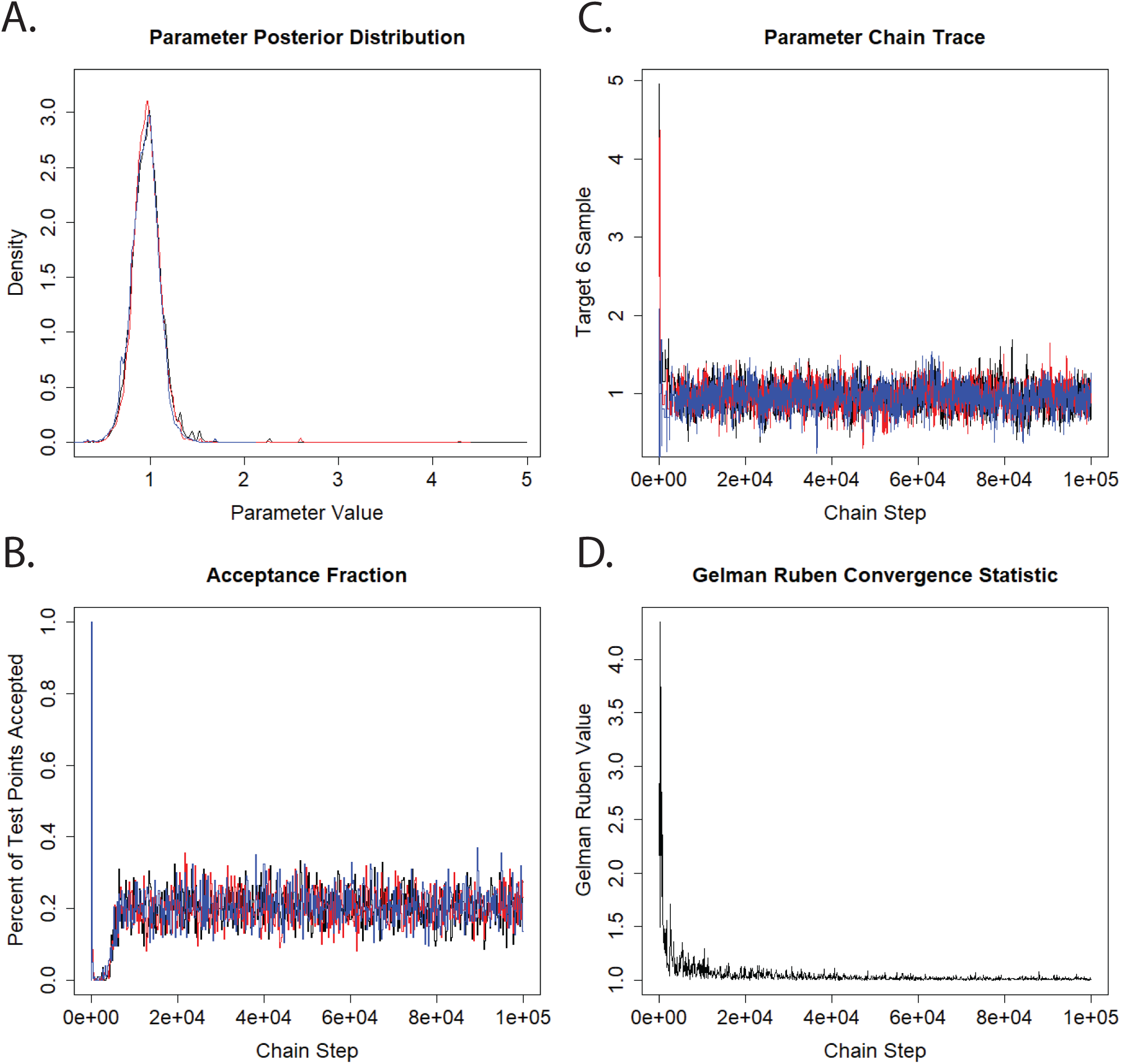
Representative images of diagnostics for Markov Chain Monte Carlo estimates of the posterior distribution of initial tumor bolus size in a single mouse. (A) The posterior distribution of the parameter for each of the three independent chains overlap, indicating agreement between the chains. (B) New steps in the Markov Chain were proposed with increased or decreased risk to achieve a desired acceptance fraction of test point of 0.20. (C) The full length traces of three independent chains (colored in black, blue, and red, respectively) with over disperse starting points. (D) The Gelman-Rubin potential scale reduction factor (PSRF) was used to assess convergence of the Markov Chains to the posterior distribution in each parameter where a value of less than 1.2 indicated that the chains have converged.

## Notes

### Competing Interest Statement

The authors have declared no competing interest.

